# FAK modulates glioblastoma stem cell energetics via regulation of glycolysis and glutamine oxidation

**DOI:** 10.1101/2024.12.11.627900

**Authors:** Roza H. A. Masalmeh, John C. Dawson, Virginia Alvarez Garcia, Morwenna T. Muir, Roderick N. Carter, Giles Hardingham, Cameron Davies, Rosina Graham, Alex von Kriegsheim, Jair Marques Junior, Chinmayi Pednekar, Steven M. Pollard, Neil O. Carragher, Valerie G. Brunton, Margaret C. Frame

**Author notes:** Correspondence (R.H.A.M.) (M.C.F.).

## Abstract

Glycolysis and the TCA cycle are reprogrammed in cancer cells to meet bioenergetic and biosynthetic demands, including by engagement with the extracellular matrix (ECM). We show that focal adhesion kinase (FAK), a mediator of integrin-ECM signalling, is driving cellular energetics in a stem cell model of glioblastoma (GBM). FAK gene deletion inhibits both glycolysis and glutamine oxidation, with increased mitochondrial fragmentation and elevated phosphorylation of the mitochondrial protein MTFR1L at S235. Simultaneously, FAK loss causes a mesenchymal to epithelial transition, enhanced acto-myosin contractility as shown by phospho-myosin light chain (p-MLC S19) and impaired cell migration/invasiveness. Rho-kinase (ROCK) inhibitors suppress p-MLC (S19) and restore glutamine oxidation and elongation of mitochondria. Thus, FAK is a key regulator of both glycolysis and glutamine oxidation mediated by acto-myosin contractility that controls both cell and mitochondrial morphology. Moreover, FAK-dependent cellular energetics are coincident properties with GBM stem cell migration, invasiveness and tumour growth in vivo.

## Introduction

Glioblastoma (GBM) is the most common and lethal form of malignant primary brain tumour ^1^. The standard treatment for this cancer involves surgical resection, radiation therapy and chemotherapy. However, none of these are curative. One of the reasons why GBM is resistant to therapy is because of the presence of a subset of cells within the tumour with high tumorigenic capacity, termed GBM stem cells^2,3^. These cells can drive recurrence and heterogeneity by rewiring their signalling and metabolism to adapt to restricted nutrition and overcome therapies^4^. The ability of cancer cells to reprogram energy metabolism has emerged as a hallmark of cancer and their ability to shift between different metabolic states has been recently recognized as a therapeutic target ^5,6^. A recent study, investigating biological traits of human GBM tumours, revealed at least four tumour cell states: proliferative/progenitor, neuronal, mitochondrial and glycolytic/plurimetabolic ^7^. GBM stem cells classified in the mitochondrial state critically depends on oxidative phosphorylation (OXPHOS) to produce energy whereas the glycolytic/plurimetabolic subtype utilises multiple energy-producing pathways, e.g., glycolysis, amino acid and lipid metabolism, granting cancer cells metabolic adaptability and protection from oxidative stress and cell death. Amongst the four subtypes, the mitochondrial subtype had significantly better clinical outcome and is more susceptible to OXPHOS inhibitors^7^. This highlights the importance of understanding metabolic pathways in GBM and the potential for finding metabolic vulnerabilities that we can exploit therapeutically.

One of the ways by which metabolism is altered in cancers is through responding to cues from the tumour microenvironment. For example, increasing extracellular matrix (ECM) stiffness can modulate glycolysis, increase the activity of metabolic enzymes, affect mitochondria morphology and function ^8–10^. These alterations help cancer cells resist oxidative stress and promote the development of invasive and metastatic phenotypes. The actin cytoskeleton can function as a scaffold for some glycolytic enzymes which can modulate glycolysis through spatial and functional regulation of these enzymes^11^. An unanswered question is whether, and how, this is connected to the role of dynamic actin remodelling in processes such as cell migration or hallmark changes in cell state, such as epithelial-to-mesenchymal transition (EMT). The links between cell-ECM mechanical cues, the actin cytoskeleton and metabolism are beginning to be identified yet we do not know which integrin may contribute.

Focal Adhesion Kinase (FAK) is a key signal transduction regulator found at integrin adhesions ^12,13^. Indeed, FAK is at the centre of one of the four focal adhesion nexuses that connect integrin complexes to actin and multiple intracellular signalling axes^13^. FAK is also known to buffer adhesion and therapeutic stress in tumour cells and promote their survival ^14^. In the context of GBM, integrin adhesion related genes (e.g., *ITGB1* and *PTK2*) are essential genetic dependencies for the injury-response transcriptional state as defined by Richards et al^15,16^.

We therefore set out to address the role of FAK in a transformed neural stem cell model of GBM that we have recently described ^17,18^ and specifically address whether FAK could link the regulation of adhesion signalling to metabolic pathways, and if so, uncover the mechanism. We found that FAK expression was increased upon oncogenic transformation and that CRISPR/Cas9-mediated genetic deletion of FAK reduced tumour growth in vivo. FAK deletion in these cells induced a profound change in both cellular morphology, and the actin cytoskeleton along with concomitant changes in gene expression consistent with transition from a mesenchymal to epithelial-like phenotype. As a result of these changes, cells were significantly less motile both in 2-dimensional culture and in 3-dimensional invasion assays. Furthermore, loss of FAK significantly suppressed rates of both glycolysis and glutamine oxidation. This was associated with enhanced actomyosin contractility especially evident at cell-cell contacts and alterations in mitochondria morphology. FAK-dependent mesenchymal morphology, spatial phospho-myosin light chain (p-MLC S19), glutamine oxidation and mitochondria morphology were restored by treatment with multiple Rho-kinase (ROCK) inhibitors. Therefore, FAK co-regulates actin, actomyosin contractility, migration and invasion via ROCK in a transformed neural stem cell GBM model and we show here that this is linked to FAK-regulated mitochondrial morphology and cellular energetics. This describes a new cellular dependency regulated by FAK in GBM cells.

## Results

### FAK controls GBM-associated cellular phenotypes

The underlying mechanisms by which FAK may contribute to GBM are poorly understood. We examined FAK’s role in GBM using a disease-relevant transformed mouse neural stem cell model that we have described previously ^17,18^. The neural stem cell model is important because neural stem cells are believed to be one of the potential cells of origin of GBM and that they give rise to GBM stem cells which are thought to drive recurrence and therapeutic resistance ^19^. We used neural stem cells which have been modified to harbour co-deletion of the genes encoding neurofibromatosis type 1 (*<u>N</u>f1*) and phosphatase and tensin homolog (*<u>P</u>ten*) and over-expression of the epidermal growth factor receptor variant III (<u>E</u>GFRvIII) (termed NPE cells). We found that the mRNA levels of *Ptk2*, the mouse gene encoding FAK, is significantly increased in transformed NPE cells when compared to untransformed parental neural stem cells (p<0.0001) (Figure 1A), and increased *Ptk2* expression is paralleled by elevated FAK protein levels in NPE cells (Figure 1B). CRISPR-Cas9-mediated targeting of *Ptk2* led, as expected, to NPE cells devoid of FAK protein (FAK -/-) (Figure 1C). We re-expressed FAK (FAK Rx) in otherwise FAK-deficient cells (FAK -/-) to levels comparable to the wild-type NPE cells to enable direct comparison of cell clones that differ only in expression of FAK (Figure 1C). To characterise the effects of deleting FAK on cellular phenotypes, we first measured cell viability of FAK Rx or FAK -/- cells cultured for 3 days and found that FAK-/- cells had significantly lower cell viability (Figure 1D). The loss of FAK also resulted in profound morphological changes. Cells expressing FAK were well attached to the cell culture substrate and displayed a spread and elongated mesenchymal-like morphology, while their comparable FAK -/- cells adopted a more rounded and less spread morphology. Moreover, FAK-deficient cells clustered tightly together with a more epithelial-like morphology and displayed reduced contact with the substratum. They displayed increased junctional F-actin at cell-cell contacts as a result of general remodelling of the actin and tubulin cytoskeletons (Figure 1E). The expression of FAK was associated with a significant enrichment of the gene set hallmark ‘epithelial to mesenchymal transition’ (FDR= 0.074) (Figure 1F). Consistent with these changes, the depletion of FAK suppressed migration speed of NPE cells in 2-dimentional (2D) culture and reduced the distance of 3-dimentional (3D) invasion into Matrigel, respectively (Figures 1G and H).

**Figure 1:**
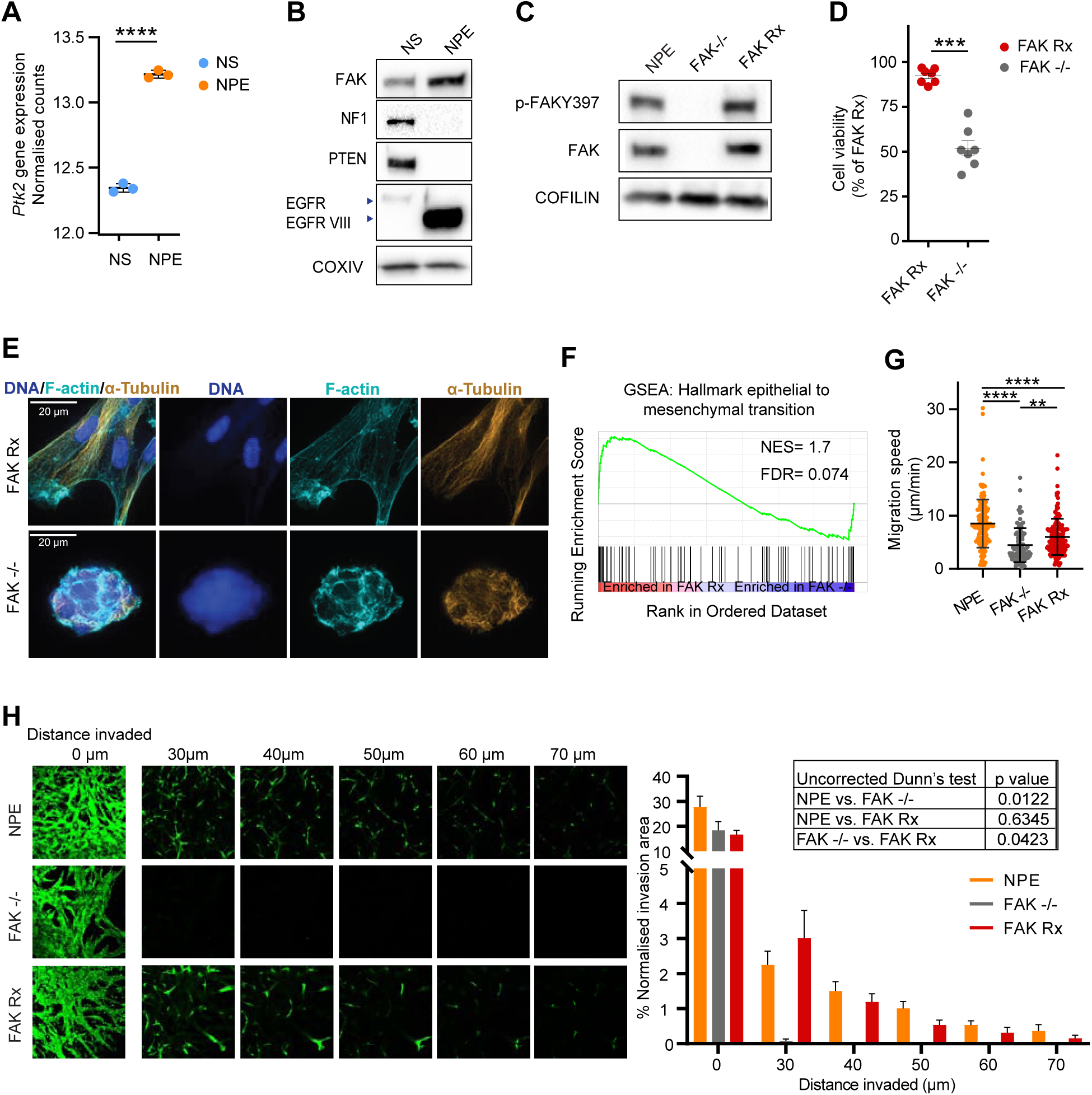
Genetic deletion of FAK drives phenotypic and transcriptional changes in a mouse model of GBM stem cells. (A) Regularised-logarithm transformation (Rlog) normalised counts of *Ptk2* in NS and NPE cells. Data obtained from (PMID: 33857425). Mean and SD are shown. Statistics: unpaired two-tailed t-test (n=3). (B) Western blot showing FAK, NF1, PTEN, EGFR, and EGFRVIII expression in NS and NPE cells. COXIV was used as a loading control. (C) Western blot analysis showing expression of p-FAK Y397 and FAK in NPE, FAK -/-, and FAK Rx cells. COFILIN was used as a loading control. (D) Cell viability of FAK Rx and FAK -/- cultured for 3 days at normal culture condition. Mean and SEM are shown. Statistics: unpaired two-tailed t-test (n=7). (E) Representative super-resolution microscopy images for FAK Rx and FAK -/- cells showing the F-actin cytoskeleton labelled with fluorophore-conjugated Phalloidin (green), α-tubulin (orange) and nuclei labelled with 40,6-diamidino-2-phenylindole (DAPI; blue). Images brightness/contrast were autoscaled individually to enable visualisation. The scale bar is 20 µm. (F) Gene set enrichment analysis (GSEA) plot of the hallmark epithelial to mesenchymal transition gene set for proteins differentially regulated in FAK Rx cells compared to FAK -/-cells (n=3). Normalised enrichment score [NES] and false discovery rate [FDR] reported. (G) Migration speed (μm/min) of NPE, FAK -/- and FAK Rx cells plated on laminin I and imaged every 10 min for 48 hrs. Each dot represents a cell, data from 2 independent cultures. Mean and SD are shown. Statistics: Kruskal-Wallis test followed by Dunn’s multiple comparison test. (H) Representative optical sections of NPE, FAK -/- and FAK Rx cells stained with Calcein AM invading through Matrigel. Right panel: the percentage of cells present in each displayed optical section of total cell area throughout the Matrigel (n=3 technical repeats). Mean and SD are shown. Statistics: Kruskal-Wallis test followed by Uncorrected Dunn’s test.

### FAK promotes glycolysis and glutamine oxidation in GBM mouse stem cells

In addition to the morphological and migratory phenotypes for which FAK is known to be crucial, we made the observation that FAK Rx culture media, containing the pH indicator phenol red, was consistently more yellow (acidic) compared to that of FAK -/- media leading us to hypothesize that FAK may also regulate cellular metabolism (Figure S1A). To test this, we performed a Seahorse XF Cell Mito Stress assay and found that FAK loss reduced both extracellular acidification rate (ECAR), a surrogate measure of glycolytic flux, and reduced oxygen consumption rate (OCR) (Figure 2A and B). This suggests that FAK loss has dampened the two main catabolic pathways glycolysis and mitochondria OXPHOS. This led FAK -/- cells to be energetically supressed and tending towards quiescence (Figure 2C). We found that spare respiration capacity, maximal respiration and respiration-linked ATP production were significantly reduced in FAK -/- cells when compared to FAK Rx cells (Figure 2D). Consistent with this, cellular metabolite profiling revealed that FAK loss reduced glycolytic intermediates, TCA cycle metabolites, ATP and its precursors (Figure 2E). In keeping with the observed metabolic effects, KEGG metabolic pathway enrichment analysis of 132 cellular metabolites supported the regulation of glycolysis, gluconeogenesis and TCA cycle in a FAK-dependent manner (Figure 2F).

**Figure 2:**
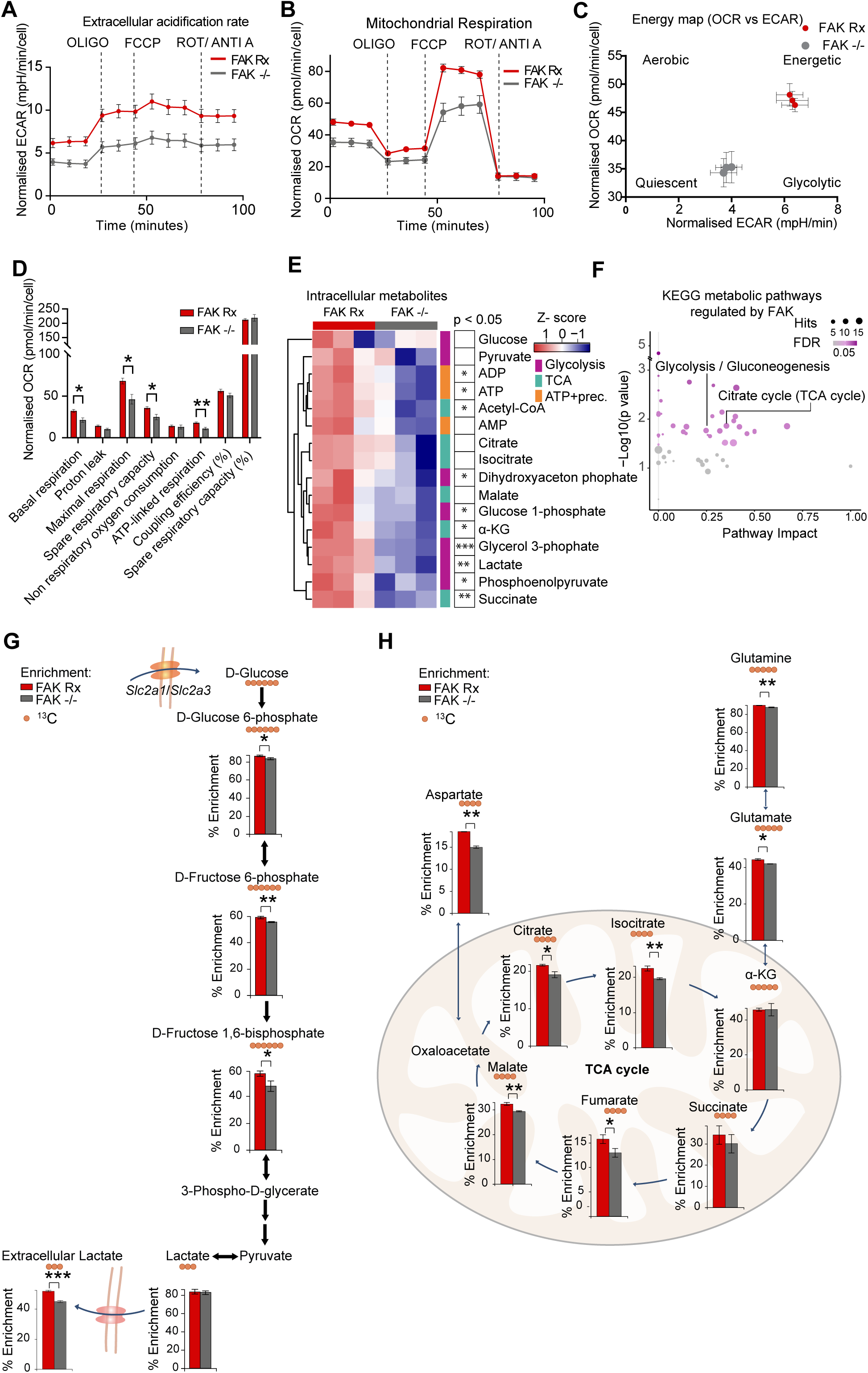
FAK promotes glycolysis, mitochondrial respiration and glutamine oxidation. Extracellular acidification rate (ECAR) (A) and oxygen consumption rate (OCR) (B) of FAK Rx and FAK -/- cells (7k cells per well, n=7 independent cultures, 3 replicate measures). Mean and SEM are shown. (C) Basal ECAR vs OCR, calculated from data in(A) and (B). Mean and SEM are shown. (D) Quantification of mitochondrial parameters calculated from the data in (B). Statistics: Mann-Whitney test. Mitochondrial stress test conditions: basal, oligomycin (OLIGO), Carbonyl cyanide-4 (trifluoromethoxy) phenylhydrazone (FCCP), a mixture of rotenone (ROT) and antimycin A (ANTI A). Mean and SEM are shown. Statics: Mann-Whitney test. (E) Heatmap of glycolysis, TCA, ATP and ATP precursors normalised peak intensity in FAK Rx cells and FAK -/- cells. Each square represents an individual replicate (n = 3 independent cultures on the same day). Statistics: unpaired two-tailed t-test (F) Bubble plot showing the glycolysis/glucogenesis and TCA cycle are among the significantly enriched KEGG pathways in pathway enrichment analysis of intracellular metabolites in FAK Rx compared to FAK -/- cells. Circles size is proportional to the number of hits in that pathway. Darker colour represents more significance. The atom fraction enrichment of glucose-derived ^13^C_6_ in glycolysis intermediates after incubation of FAK Rx and FAK -/- cells with ^13^C_6_ D-glucose for 1 h (G) or ^13^C_5_ glutamine for 3h (H) (n = 3 independent cultures on the same day). The main isotopologue of each metabolite is shown and plotted as the fraction of the sum of all isotopologues. Mean and SD are shown. Statistics: unpaired two-tailed t-test.

To further characterise FAK’s role in regulating glycolysis and the TCA cycle, we performed stable isotope-labelled ^13^C_6_-glucose tracing in FAK-expressing and FAK-deficient cells. We found significantly reduced labelling of glycolytic intermediates within cells and of secreted lactate, the end product of anaerobic glycolysis, in the absence of FAK (Figure 2G). We also traced ^13^C_5_-glutamine and found reduced labelling in most TCA cycle metabolites in FAK -/- cells compared to FAK Rx cells (Figure 2H), indicative of a role for FAK in efficient glutamine oxidation. Taken together, these findings have uncovered a previously unknown role for FAK in promoting both efficient glycolysis and glutamine oxidation.

We noted that FAK’s role in promoting glycolysis was consistently associated with higher levels of glucose transporter proteins, namely Glucose transporter 1 (GLUT1, encoded by the *Slc2a1* gene) and Glucose transporter 3 (GLUT 3, encoded by *Slc2a3* gene) (Figure S1B and C), although we have not addressed causality of these changes.

### FAK regulates the length of mitochondria

Previous studies have suggested that the shape of mitochondria is linked to their function, and is responsive to nutrients and energy demands (reviewed in ^20^). Specifically, it is proposed that elongated mitochondria are associated with efficient OXPHOS ^21–23^. Given this, and FAK’s role in regulating cell adhesion and cytoskeletal structure in the NPE model of GBM we have used here, we asked whether FAK’s regulation of glutamine oxidation may be functionally linked to FAK-dependent control of mitochondria morphology. We used Spinning disk super-resolution by Optical pixel Reassignment (SoRa) microscopy to image the mitochondrial network stained with MitoTracker (Figure 3A). We then used mitochondria features extraction tool (called Mitochondria Analyser Fiji/ImageJ plugin^24,25^) to define the mitochondrial network and extract different mitochondria features. This revealed that FAK’s loss decreased the mitochondria mean branch length (Figure 3B), implying that FAK promotes elongated and filamentous mitochondria; a morphology associated with more efficient OXPHOS (Figure 2B, D and H). A comprehensive phosphoproteomic analysis of FAK-expressing and FAK-deficient cells revealed a number of changes, including that deletion of FAK leads to a significant increase in phosphorylated MTFR1L (p-MTFR1L S235) (Figure 3C, and D). This modification, which is the orthologous site of S238 in human MTFR1L and is phosphorylated by AMPK, is known to result in mitochondrial fragmentation under metabolic stress ^26^ and is therefore a candidate mediator of the effects of FAK loss on mitochondria morphology and GBM cellular energy production.

**Figure 3:**
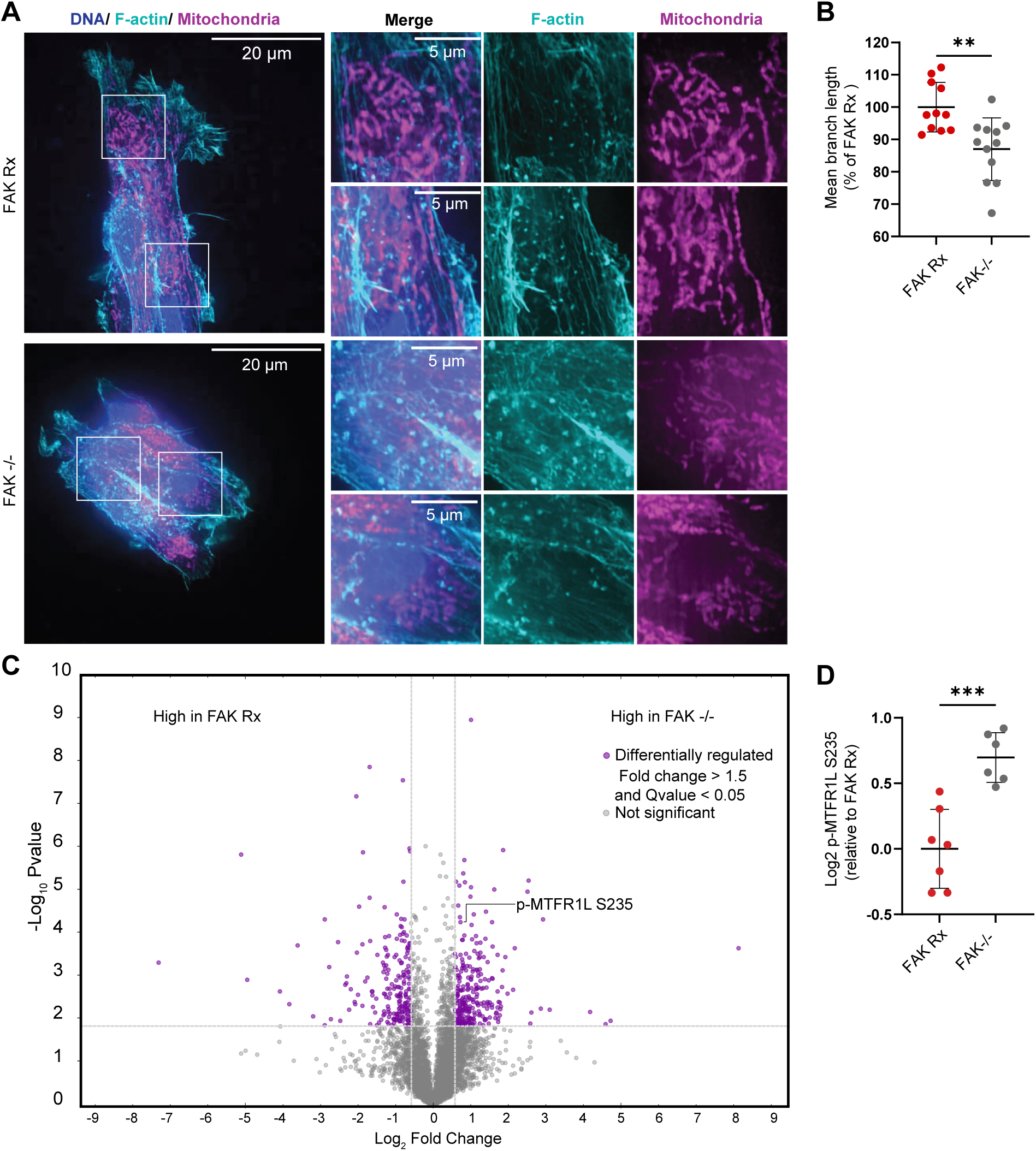
FAK is associated with elongated mitochondria. (A) Representative super-resolution microscopy images for FAK Rx and FAK -/- cells showing mitochondrial morphology stained with MitoTracker Deep Red FM (magenta), the F-actin cytoskeleton labelled with fluorophore-conjugated phalloidin (cyan), and nuclei labelled with DAPI (blue). Scale bar is 20 µm in the full field of view and 5 µm in insets. (B) Quantification of mitochondrial mean fragment length in FAK Rx and FAK -/- cells. n = 2 independent experiments. Each dot represents a field of view. Mean and SD are shown. Statistics: unpaired two-tailed t-test. (C) Volcano plot of post-translational modifications highlighting the differentially modified proteins in red. p-MTFR1L S235 is labelled. n=3 independent cultures. (D) p-MTFR1L S235 abundance in FAK Rx and FAK -/- cells. n=7 (FAK Rx) and n=6 (FAK null) independent cultures. Data from two separate experiments conducted on different days are shown. Mean and SEM are shown. Statistics: unpaired two-tailed t-test.

### FAK regulates length of mitochondria via ROCK

The shape and function of mitochondria is responsive to mechanical signals stimulated by interaction with the ECM ^8,10^. FAK is implicated in mechanotransduction by connecting signals from the integrin family of ECM adhesion receptors to the actin cytoskeleton and actomyosin contractility^27^, and this has been shown to regulate the switch between epithelial- and mesenchymal-like colon cancer cells^28^. Since the mesenchymal-like phenotype of the transformed NPE cells we used here was also FAK-dependent (Figure 1E and F), we next addressed whether FAK regulates mechanotransduction in NPE cells and, if so, whether this is associated with changes in mitochondrial morphology and metabolism.

Phosphorylation of the regulatory light chain of myosin type II at serine(S)19 (hereafter p-MLC(S19)), initiates actin-myosin interactions and assembles myosin filaments driving actomyosin contractility^29^. We therefore used pMLC2(S19) as a surrogate measure of cellular actomyosin contractility. In order to localise pMLC2(S19) at a subcellular level in FAK-expressing and FAK-deficient cells, we used SoRa super-resolution microscopy and quantified pMLC2(S19) intensity at cell-cell contacts, cell edge and cytoplasmic locales. We found that FAK -/- cells had higher mean intensity of pMLC2(S19) at all of these subcellular locales, while noting that pMLC2(S19) mean intensity was the highest at cell-cell contacts with intense junctional F-actin (Figure 4A and B). This implied that the increased cell-cell contacts formed as a result of FAK depletion were associated with highly localised F-actin and acto-myosin contractility at the intercellular junctions. Next, we used inhibitors of ROCK, the kinase the maintains MLC2(S19) phosphorylation^30^. We treated FAK -/- cells with 2 μM of the ROCK inhibitor GSK269962A which reduced pMLC2(S19) signal across the cells and induced cell spreading, suppressing cell-cell contacts and restoring a mesenchymal-like morphology (Figure 4A and B). Taken together, these data demonstrate that pMLC2(S19), and so by inference actomyosin contractility, concentrates at cell-cell contacts in FAK-deficient cells and that treating the cells with a ROCK inhibitor suppresses cell-cell contacts, enhances cell spreading and restoration of the mesenchymal-like morphology typical of FAK-expressing NPE cells.

**Figure 4:**
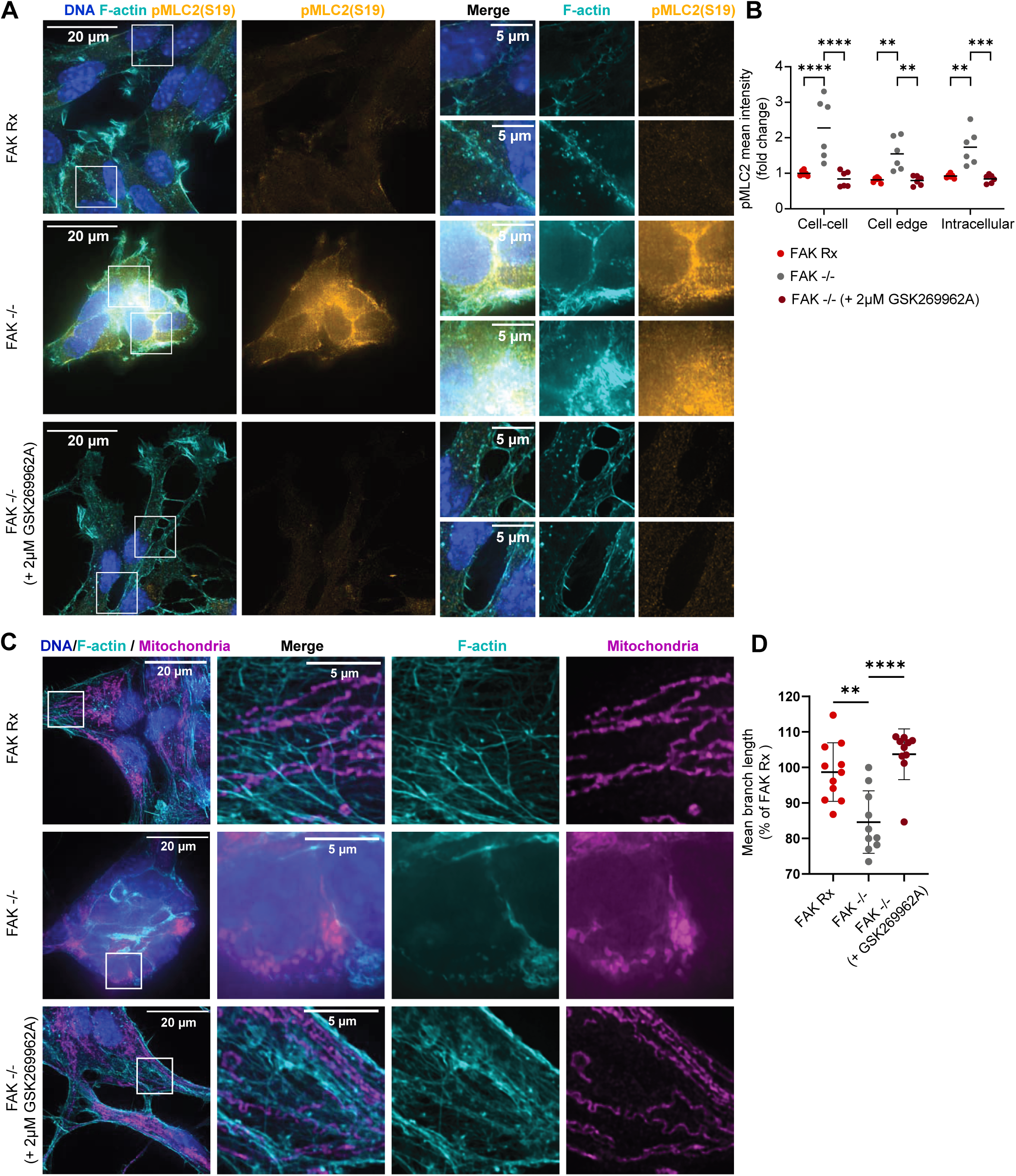
FAK regulates mitochondria morphology through ROCK-pMLC2 S(19) pathway. (A) Representative super-resolution microscopy images for FAK Rx, FAK -/- cells treated with or without 2 µM GSK269962A showing pMLC2 (S19) (yellow), the actin cytoskeleton labelled with fluorophore-conjugated Phalloidin (cyan) and nuclei labelled with DAPI (blue). Cell-cell contacts are shown in insets. Scale bar is 20 µm in the full field of view and 5 µm in insets. (B) Quantification of pMLC2 (S19) mean fluorescence intensity at cell-cell contacts, cell edge and intracellularly. n = 2 independent experiments. Each dot represents a field of view. Mean and SD are shown. Statistics: two way ANOVA followed by Tukey’s multiple comparisons test. (C) Representative super-resolution microscopy images for FAK Rx, FAK -/- cells treated with or without 2 µM GSK269962A showing mitochondrial morphology stained with MitoTracker Deep Red FM (magenta), the F-actin cytoskeleton labelled with fluorophore-conjugated phalloidin (cyan), and nuclei labelled with DAPI (blue). Scale bar is 20 μm in the full field of view and 5 μm in insets. (D) Quantification of mitochondrial mean fragment length in FAK Rx and FAK -/- cells with or without 2 μM GSK269962A. n = 2 independent experiments. Each dot represents a field of view. Mean and SD are shown. Statistics: one way ANOVA followed by Tukey’s multiple comparisons test.

Next, to address whether FAK’s regulation of mitochondrial morphology may also be dependent on its role in mechanotransduction, we used SoRa super-resolution microscopy to characterise mitochondria morphology. We found that treating FAK-deficient cells with ROCK inhibitors GSK269962A (2 µM) or Y27632 (20 µM) restored mitochondria morphology to that exhibited by FAK-expressing cells, as judged by increased mitochondrial mean branch length (Figure 4C and D and Figure S2A and B). This implies that inhibiting ROCK-pMLC(S19) signalling enhances mitochondria elongation, linking FAK-mediated actomyosin contractility to mitochondrial morphology.

### ROCK inhibitors increase glutamine oxidation and cell viability in FAK -/- cells

We hypothesized that the observed restitution of mitochondrial morphology might be associated with the restoration of normal glutamine oxidation found by re-expressing FAK in otherwise FAK-deficient cells. To test this, we analysed glutamine flux by performing a ^13^C_5_ labelled glutamine tracer analysis of the TCA cycle metabolites and found that both ROCK inhibitors GSK269962A (2 µM) and Y27632 (20 µM) increased glutamine oxidation to normal levels (Figure 5A and Figure S3A). Together, these data imply that FAK regulates mitochondrial morphology and glutamine oxidation by counteracting the ROCK-pMLC2(S19) signalling axis. We found that restoration of glutamine oxidation by ROCK inhibitor treatment rescued the loss of viability we observed in FAK-deficient cells (Figure 1D; Figure 5B). These data indicate that FAK mediated suppression of the ROCK/p-MLC2(S19) pathway and actomyosin contractility not only control maintenance of the mesenchymal-like morphology and migration/invasion, but are also inextricably linked to mitochondrial morphology, cell energy production and cell viability.

**Figure 5:**
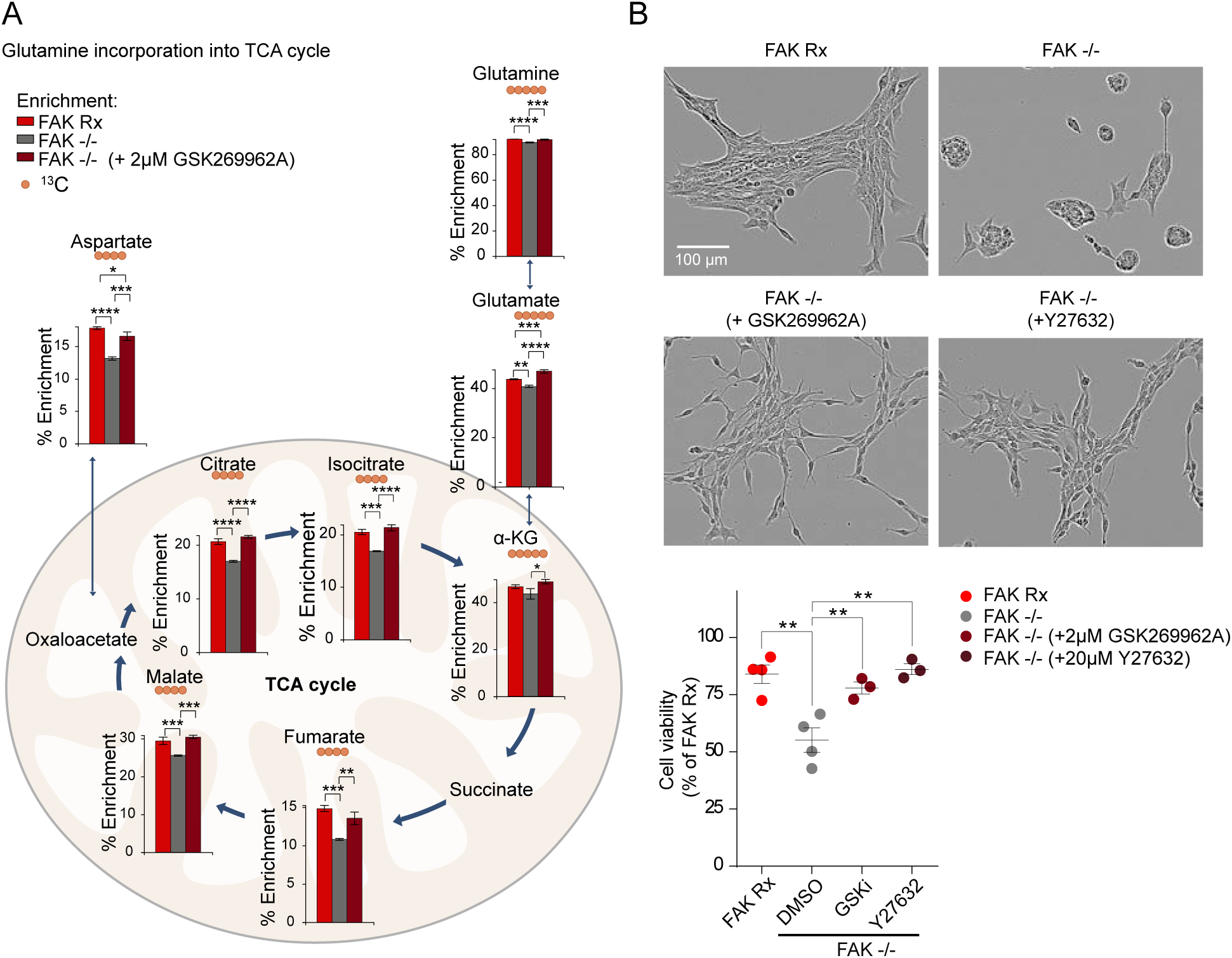
Treating FAK -/- cells with ROCK inhibitor GSK269962A increases glutamine oxidation and cell viability. (A) The atom fraction enrichment of glutamine-derived ^13^C in TCA intermediates in FAK Rx, FAK -/- and FAK -/- cells treated with or without 2 µM GSK269962A after 3 h incubation with ^13^C_5_ glutamine supplemented medium. The main isotopologue of each metabolite is shown and plotted as the fraction of the sum of all isotopologues. Mean and SD are shown. Statistics: one-way ANOVA with a Tukey multiple test correction (n = 3 independent cultures on the same day). (B) Top panel, representative phase-contrast images of FAK Rx, FAK -/- treated with and without ROCK inhibitors (+2μM GSK269962A and +20μM Y27632) for 3 days. Cell viability of FAK Rx, FAK -/- treated with and without ROCK inhibitors (+2μM GSK269962A and 20μM Y27632) for 3 days. Mean and SEM are shown. Statistics: one-way ANOVA with a Dunnett multiple test correction (n=>3 independent experiments).

### FAK controls invasion, proliferation and survival in vivo

Loss of FAK led to a significant decline in key hallmarks of the malignant phenotype in the transformed neural stem cell model of GBM we used here, causing them to exhibit reduced invasion, migration, cell viability and suppressed glycolysis and glutamine oxidation rendering cells less energetic in vitro. To investigate whether this would translate to beneficial properties in vivo, we performed an intracranial transplantation of GFP and luciferase-expressing FAK -/- cells or FAK-Rx cells into the right-side brain hemisphere of CD1 nude mice. We found that mice injected with cells lacking FAK had a significantly increased overall survival (Figure 6A). CD1-nude mice with intracranial tumours formed after injecting FAK Rx cells showed higher total body flux of the luciferase reporter when compared to FAK -/- tumours at day 7 and day 17 (Figure 5B), suggesting that FAK expression increases tumour growth. This was associated with a higher percentage of Ki67-expressing cells in sections of FAK-expressing intracranial tumours at 17 days (Figure 6C and D). Moreover, FAK loss significantly reduced the distance NPE cells could invade into the healthy brain tissue from the tumour margin, a measure of their in vivo invasive capacity, implying a role for FAK in cancer cells invasion in vivo (Figure 6E).

**Figure 6:**
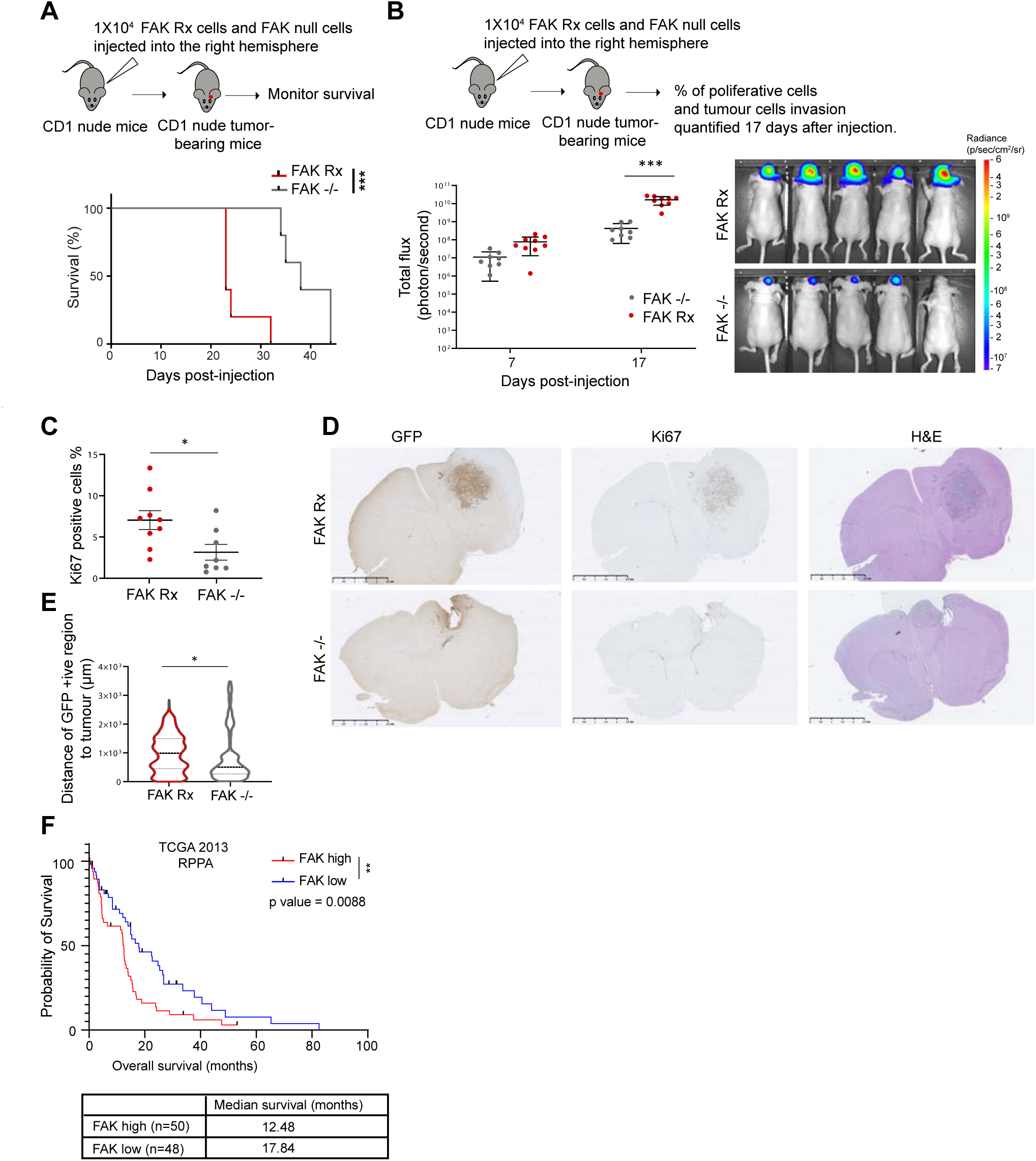
FAK protein correlates with worse overall survival in vivo and in GBM patients. (A) Kaplan-Meier curve of CD1-nude mice bearing intracranial tumours from injecting FAK Rx and FAK -/- cells. Statistics: Mantel-Cox test, n=5 for each group. (B) Total body flux of CD1-nude mice bearing intracranial tumours from injecting luciferase-expressing FAK Rx (n=9 mice) and FAK null cells (n=8 mice) at 7 and 17 days after injection measured by IVIS imaging. Mean and SEM are shown. Statistics: 2-way ANOVA with Tukey post hoc adjustment for multiple comparisons. Right panel: representative in vivo bioluminescent images of CD1-nude mice bearing tumours from injecting FAK Rx and FAK null cells at day 17. The heat map superimposed over the mouse heads represents the radiance (photons/s/cm2/sr). (C) Percentage of KI67 + cells in sections of intracranial tumours from injecting FAK Rx (n=9 mice) and FAK null cells (n=8 mice) at 17 days after injection. Mean and SEM are shown. Statistics: unpaired two-tailed t-test. (D) Distance of GFP+ tumour cells to tumour edge in sections of intracranial tumours from injecting FAK Rx (n=4) and FAK null cells (n=4) at day 17 after injection. Mean and SEM are shown. Statistics: unpaired two-tailed t-test. (E) Representative images of the immunohistochemical (IHC) staining for GFP, Ki67 and H&E stained intracranial tumours from

Finally, to investigate the relevance of our findings to human GBM, we used data from the Cancer Genome Atlas (TCGA) to examine whether there was an association between FAK protein levels (measured by Reverse Phase Protein Array) and patient survival in adult GBM cases. We identified a group of patients with high (n=50) or low (n=48) FAK expression, representing those with higher and lower quartile FAK expression, respectively. Our analysis revealed an inverse correlation between FAK protein levels and overall patient survival (p = 0.0088), consistent with FAK contributing to GBM pathogenesis in patients (Figure 6F).

## Discussion

Metabolic processes play critical roles in GBM as highlighted by the recently identified metabolic subtypes with associated metabolic vulnerabilities^7,31^. Moreover, many GBM driver mutations, such as loss of the tumour suppressor *PTEN*, or amplification of the genes for receptor tyrosine kinase such as *EGFR* or platelet-derived growth factor receptor A (*PDGFRA*) can enhance PI3K-AKT-mTOR signalling that, in turn, confers metabolic plasticity, enabling GBM to tailor different metabolic pathways to their requirements ^32^. Aerobic glycolysis has been established as a metabolic pathway beneficial for tumour growth, and more recently the TCA cycle has also been shown as important for cancer cells both in mouse models and patients ^33,34^. The role of the ECM in modulating these pathways to enhance metabolic plasticity and resilience, including under therapeutic and mechanical stress, is beginning to be elucidated and is cell-type and tissue specific. ^8,10,35,36^. For example, mechanical cues from the ECM influence ATP production and recycling, aiding the invasive migration of pancreatic cancer cells^10^. In addition, increased stiffness boosts glycolysis in human bronchial epithelial cells (HBECs), while transformed non-small cell lung cancer cells maintain high glycolytic rates despite changes in environmental mechanics ^9^, demonstrating cell- and tumour-context dependency. Furthermore, cell-ECM-mediated mechano-sensing and downstream signalling are also known to regulate mitochondrial structure and function, conferring resilience to redox stress in malignant human mammary epithelial cells^8^.

Despite these advances, how specific adhesion proteins that function downstream of ECM-cell interactions influence cellular energetics and metabolic flexibility is still unclear. The work we present here has uncovered a novel link by which a pivotal adhesion protein, namely FAK, is vital for optimal throughput of both glycolysis and the glutamine oxidation to ensure transformed neural stem cells have sufficient energy production. When the gene encoding FAK is deleted, the cellular energy production is significantly impaired and glutamine oxidation is supressed. This is mediated by regulation of actomyosin contractility, as evidenced by p-MLS-S19), likely via ROCK, including at cell-cell junctional contacts in FAK depleted cells induced to undergo a mesenchymal-like to epithelial-like morphological transition. This is important because glutamine is the major molecule that replenishes the TCA cycle in glioma cell lines and its uptake is highly enhanced with minimal uptake in the surrounding brain tissue in both mice and in patients ^37^.

In addition to the suppression of cellular energetics as a result of impaired glycolysis and glutamine oxidation in FAK depleted transformed neural stem cells, we found visible changes to mitochondrial morphology. Mitochondrial function is known to be closely connected to their structure, which can have impact in human health and disease in a context dependent manner ^38^. We found that FAK’s role in promoting glutamine oxidation was associated with an elongated mitochondrial network. The changes in mitochondria morphology when FAK was depleted was dependent on FAK-mediated suppression of signalling to p-MLC2. Indeed, treating FAK-deficient cells with two distinct ROCK inhibitors (GSK269962A and Y27632) restored mitochondria morphology to the elongated form that was associated with increased glutamine incorporation into TCA cycle. In keeping with this, RhoA activity has been shown to promote mitochondrial fission in cardiomyocytes by phosphorylating the protein Drp1 at serine-616, leading to its re-localization to mitochondria^39^, and this is blocked by treatment with the ROCK inhibitor Y-27632 ^39^. Additionally, higher Myosin II activity is associated with mitochondria fragmentation in human melanoma cells ^36^. We have concluded that one of FAK’s key roles in the GBM stem cell model is to sustain cellular energetics via both glycolysis and glutamine oxidation, linked to mitochondrial structural changes that are accompanied by phosphorylation of the protein MTFR1L on S235 that is recognised to promote mitochondrial dynamics^26^. We propose this contributes to the impaired tumour growth and overall survival in vivo when cells are implanted into brains of recipient mice.

As mentioned, the work we report here used a mesenchymal-like GBM stem cell model; when we convert these into more epithelial-like cells by deletion of the gene encoding FAK, both glycolysis and mitochondrial respiration are significantly impaired. However, previous studies in a pancreatic ductal adenocarcinoma (PDAC) epithelial cell line and in epithelial-like GBM cell lines, both cultured in serum-containing media, rather than the serum-free stem cell medium we used here, suggests that FAK promotes glycolysis but suppresses mitochondrial respiration under these conditions^40,41^. Taken together, FAK’s role in modulating metabolism in epithelial-like cells differs from its role in mesenchymal-like cells of the same origin. In this regard, we note that genome-wide CRISPR screens identified FAK as a fitness gene in the transcriptionally-defined ‘injury response’ GBM stem cell subtype but not in those defined as the ‘developmental’ GBM stem cell subtype ^15,16^. Injury response GBM stem cells map to the mesenchymal-like and glycolytic/plurimetabolic cell states, while the developmental GBM stem cells, which are much less dependent on integrin signalling and FAK, map to non-mesenchymal-like cell states, such as ‘neural precursor’, ‘astrocytic’ and ‘oligodendrocyte progenitor’-like^15,16^. Our data imply that mesenchymal-like cells are dependent on FAK for energy production.

Our findings offer one possible explanation for why the genes encoding FAK, and other integrin or their effector adhesion proteins, are amongst the primary ‘fitness genes’ identified in drop-out screens in the highly treatment-resistant mesenchymal-like subtypes/GBM cell states^15,16^. The novel role we have identified for FAK specifically in promoting metabolic pathways in mesenchymal-like cells implies that FAK controls the bioenergetic and biosynthetic requirements for the metabolic flexibility that characterises these GBM tumours. In turn, this likely contributes to their growth in vivo and resistance to multiple therapeutics^42^. Our data imply that targeting FAK is likely to limit cellular energy production and metabolic flexibility of mesenchymal-like and glycolytic/plurimetabolic GBM tumours. We therefore suggest that a combination of targeting FAK together with agents that inhibit residual metabolic pathways in susceptible GBM subtypes may lead to enhanced therapeutic benefit.

## Methods

### Experimental model and subject details

#### Cell culture

NPE cells were generated using CRISPR/Cas9 mutagenesis from neural stem cells of adult male C57BL/6-SCRM mice, as previously described^17^. NPE cells were maintained in 5% CO_2_ at 37 °C and were routinely tested for mycoplasma. Cell numbers were obtained using the Countess automated cell counter (Thermo Fisher Scientific) by Trypan blue exclusion. NPE cells were cultured in Dulbecco’s Modified Eagle Medium and Ham’s Nutrient Mixture F12 (DMEM/F12; Sigma-Aldrich, #D8437) supplemented with 1.45 g/L D-Glucose (Sigma-Aldrich, #G8644), 120 μg/mL bovine serum albumin (BSA) fraction V solution (Gibco, #15260-037), 100 μM β-mercaptoethanol (Gibco; #31350-010), 1X MEM non-essential amino acid (MEM-NEAA) solution (Gibco; #11140-035), 0.5X B-27 (Gibco; #17504-044) supplement, 0.5X N-2 supplement (Gibco; #15140-122), 10 ng/mL murine EGF (Peprotech; #315-09), 10 ng/mL human b-FGF (Peprotech; #100-18b), and 1 μg/mL laminin-I (R&D Systems; #3446-005-01). Cells were passaged every second day by dissociation with Accutase (Sigma-Aldrich; #A6964).

#### In vivo experiments

All treatments and procedures with mice were performed in accordance with protocols approved by Home Office UK guidelines in a designated facility under a project license PP7510272 held by V.B. at the University of Edinburgh. Up to 6 mice were caged in ventilated cages in a pathogen-free facility on a 12-hour light/dark cycle with unlimited access to food and water. 6–15-week-old female CD-1 nude mice (Charles River, Strain number 086) were injected intracranially with 10,000 cells of either NPE-derived FAK-/- cells or NPE-derived FAK Rx cells suspended in 2μl neural stem cell medium following administration of isoflurane general anaesthesia. Mice were allocated to groups randomly.

Mice were given analgesic buprenorphine subcutaneously during surgery and the analgesic carprofen was administered in drinking water for 48 hours post-surgery. Experimental mice were placed in a 25-30°C heat box during recovery from anaesthesia. Intracranial injections were performed in the mouse striatum at coordinates 0.6 mm anterior and 1.5 mm lateral to the bregma and 2.4 mm deep. Cells were prepared as previously described^43^. Bioluminescent imaging was performed while mice were under anaesthesia twice weekly by subcutaneous injection of 150 mg/kg luciferase using an IVIS Lumina S5 system (Revvity) for monitoring tumour growth. For survival experiments, mice were sacrificed by cervical dislocation upon the development of symptoms indicating the presence of a brain tumour such as lethargy, instability, hunched posture and weight loss or until a maximum time-point of 90 days post-injection. For endpoint experiments, mice were sacrificed by cervical dislocation 17 days after cell implantation. Brains were collected immediately after sacrifice and fixed in 10% Neutral Buffered Formalin (Merck; HT501128-4L) overnight.

## Method details

### Cell Treatments

NPE cells were transfected by electroporation using a Lonza® Nucleofector™ 2b device and the Lonza® Mouse Neural Stem Cell Nucleofector™ Kit (Lonza; #VPG-1004) according to the manufacturer’s instructions using the T-030 pulse code.

### Plasmids

Plasmids for CRISPR/Cas9-mediated *Ptk2* knockout were generated by cloning oligonucleotides encoding 1 sgRNAs targeting the *Ptk2* gene (Table S1) between the BbsI sites of the pSpCas9(BB)-2A-GFP (PX458) plasmid ^44^ (a gift from Feng Zhang (Addgene plasmid # 48138; RRID: Addgene_48138)). For *Ptk2* re-expression, total RNA was extracted from NPE cells and reverse-transcribed to produce a cDNA library according to the manufacturer’s protocols (Qiagen; #74104 and Thermo Scientific; #K1621). The *Ptk2* gene was amplified from the cDNA library using specific primers containing Kozak sequence^45^ before the start codon with 15 bp overhangs (L_*Ptk2*_HD and R_*Ptk2*_HD; Table S1) to facilitate In-Fusion cloning (TakaraBio) into pQCXIN plasmid (a gift from Toby Hurd, Institute of Genetics and Cancer, The University of Edinburgh) by In-Fusion cloning following the manufacturer’s instructions.

### CRISPR/Cas9 genome editing

1 x 10^6^ NPE cells were electroporated as described with 5 μg of either pSpCas9(BB)-2A-GFP (PX458) with *Ptk2* sgRNA (sgRNA: 5’-GCAGTAGTGAGCCAACCACCT). This process was repeated twice at 48 h intervals. Cells were suspended in 5% BSA (Merck; #12659) in PBS and single cells were sorted using a BD FACSJazz™ cell sorter into wells of a 96-well cell culture plate. Cells were incubated in a humidified 37°C incubator at 5% CO_2_ until visible colonies were formed. Colonies were passaged into two wells of a 6-well plate, one of which was lysed directly in Laemmi buffer (50 mM Tris-HCl pH 6.8, 10% glycerol, 5% SDS, 5% β-mercaptoethanol, bromophenol blue), and processed for western blotting to screen for FAK depletion. Where FAK loss was detected, the remaining cells were passaged for further use.

### DNA construct expression

For re-expression of the *Ptk2* gene in NPE FAK -/- cells, 2 x 10^6^ cells were electroporated with 8 μg of FAK-pQCXIN. 48 h later, FAK-expressing cells were selected by addition of 0.5 mg/mL G418 (Merck; #A1720). G418 selection was continued for 2 weeks to select only cells that had stably integrated the FAK-pQCXIN construct. Expression of FAK in G418-selected cells was confirmed by Western blotting. Selection was repeated every 3 months to ensure that the population retained FAK expression.

### Cell lysis and western blotting

Cells were washed once with cold PBS and lysed in cold RIPA buffer (150 mM NaCl, 50 mM Tris-HCl pH 8, 1% v/v Triton X-100, 0.5% w/v sodium deoxycholate, 0.1% w/v sodium dodecyl sulphate) supplemented with cOmplete™ ULTRA Protease Inhibitor (Roche; #5892953001) and PhosSTOP™ Phosphatase Inhibitor (Roche; #4906845001) cocktail tablets for 15 minutes at 4 °C with gentle agitation. Crude extracts were collected into microcentrifuge tubes and centrifuged at 4 °C for 15 minutes at 19,000 x *g*. Protein content was quantified using the Pierce™ BCA assay kit (Thermo Scientific; #23225) according to the manufacturer’s protocol. 20 μg protein was diluted in Invitrogen novex NuPAGE LDS Sample Buffer (4X) and heated to 95 °C for 7 minutes. Proteins were resolved by SDS-polyacrylamide gel electrophoresis using 4-15% Mini-PROTEAN® TGX™ gels (BioRad; #4561086) and transferred to PVDF membranes (BioRad; #1704157) using the Trans-Blot Turbo semi-dry transfer system. Membranes were blocked by incubation with Tris-buffered saline (TBS) supplemented with 5% BSA and 0.1% TWEEN® 20 (Millipore; #11332465001) (TBS-T) for 1 hour at room temperature with gentle agitation. Primary antibodies were diluted in TBS-T supplemented with 5% BSA and incubated with membranes overnight at 4 °C with gentle agitation. Membranes were washed 3 times with TBS-T at room temperature for 15 minutes with agitation. Secondary antibodies were diluted in TBS-T supplemented with 5% BSA and incubated with membranes for 45 min at room temperature with agitation. Membranes were washed a further 3 times in TBS-T and bound antibodies were visualised using the Clarity Western ECL substrate (BioRad; #1705061) with a ChemiDoc MP system (BioRad). Images were collected and quantified using the BioRad ImageLab software (v6.1).

### In vitro 2D migration assay

500 cells/well were plated into an IncuCyte® ImageLock 96-well plate (Sartorius; #BA-04856) and imaged using a 10X objective every 10 min for a period of 48 h. Single-cell tracking was performed in Fiji/ImageJ (v2.14.0/1.54f)^25^ using Trackmate7 plugin^46^.

### In vitro respirometry

7000 cells/well were seeded in a Sea horse XF24 cell culture microplate for 48 hours (Part # 102342-100) after which medium was replaced with Seahorse XF medium (#103680-100) supplemented with 10mM Glucose (103577-100), 2mM Glutamine (103579-100), 1mM pyruvate, 10 ng/mL murine EGF, and 10 ng/mL human b-FGF and incubated in a CO_2_ free incubator for 30 minutes before the assay. During respirometry, the following inhibitors were sequentially added via injection ports; oligomycin (1μM final), FCCP (2μM final), and antimycin A/rotenone (0.5 μM final) during (concentrated stock solutions solubilized in Seahorse XF medium (#103680-100) for mitochondrial stress test compounds). XFe24 sensor cartridges containing 1ml Seahorse XF Calibrant Solution/well were incubated for 16 hrs before the assay in a CO2 free incubator. Measurements were performed using Seahorse XFe24 Analyzer (Agilent) and data visualised and pre-processed using the Wave software (v.2.6.3.5). Cells were dissociated with Accutase (Sigma-Aldrich; #A6964) and counted after the assay. Data was normalised to cell number. ‘Non respiratory OCR’ was defined as measurement 10 (ie the OCR remaining following complete respiratory block with antimycin/rotenone). ‘Basal respiration’ was calculated by subtracting OCR at measurement 10, from OCR at measurement 3 (the basal OCR). Other parameters were similarly derived by subtraction; ‘Proton Leak’ measurement 10 from 4, Maximal respiration measurement 10 from 7, ‘Spare respiratory capacity’ measurement 3 from 7, ATP-linked respiration (ATP production) measurement 4 from 3.

### Drug treatments

For drug treatment, cells were incubated with the indicated doses of either Y-27632 dihydrochloride (Tocris; #1254), GSK269962A (Cayman Chemical; #19180), or 0.04% DMSO. Drugs were solubilised in DMSO to a stock concentration of 50 mM and10 mM, respectively.

### LC-MS metabolomics

500,000 cells/well were seeded in a 6-well plate and cultured for 48 hours before metabolites were extracted with a solvent mixture (50% methanol, 30% acetonitrile, 20% water v/v) on dry ice for 15 minutes with agitation. The extraction solution was then centrifuged at 16,100 x g for 10 minutes at 4°C. The supernatants stored at −75°C prior to LC-MS analysis. Plates were left in fume hood to dry and protein quantification was performed using the Lowry assay.

Samples were randomised and metabolites were analysed on a Dionex Ultimate 3000 UHPLC (Thermo Fisher Scientific) coupled to a Q Exactive Hybrid Orbitrap MS (Thermo Fisher Scientific). Hydrophilic interaction liquid chromatography (HILIC) using a ZIC-pHILIC analytical column (2.1 × 150 mm, SeQuant-MerckMillipore) coupled with a guard column (2.1 × 20 mm) was used for chromatographic separation of metabolites. A gradient using mobile phase A (20 mM ammonium carbonate, 0.01% (v/v) ammonium hydroxide), and B (acetonitrile) was used. The gradient was set from 95% to 5% B over 20 min and reequilibrated to initial conditions for 7 min. The flow rate was 200 ml/min, and the temperature was at 45°C. The injection volume was 5 µl. MS data was acquired in positive/negative polarity switch mode in the m/z range of 70–900 Da, with a resolving power of 70,000 (FWHM). Metabolites were identified by matching accurate mass measurements (< 5 ppm) and retention times to an in-house library of standards. Skyline 21.2.0.568^47^ was used to quantify the metabolites by integrating the area under the curve and normalizing the values to total protein content. Metaboanalyst 5.0^48^ was used to perform log transformation (base 10), auto scaling (data mean-centred and divided by the standard deviation of each variable) and KEGG metabolic pathway enrichment analysis.

### ^13^C_6_-glucose and ^13^C_5_-glutamine LC-MS metabolomics

500,000 cells/well were seeded in a 6-well plate and cultured for 48 hours before media was exchanged with media containing ^13^C_6_-Glucose (Sigma Aldrich;389374) at 4.5 g/L and incubated at 37°C for 1 hr for ^13^C_6_-glucose tracing. For glutamine tracing media was replaced with media containing ^13^C_5_-glutamine (Sigma Aldrich; 605166) at 0.365 g/L and incubated at 37°C for 3 hr. Cells were washed twice with PBS and metabolites were extracted with a solvent mixture (50% methanol, 30% acetonitrile, and 20% water (v/v)) on dry ice for 15 minutes with agitation. The extraction solution was then centrifuged at 16,100 x g for 10 minutes at 4°C. The supernatants were stored at −75°C prior to LC-MS analysis. Metabolites were analysed as described above. Isotope natural abundance was corrected and % Enrichment of isotopolouges was calculated using “accucor” package in R.

### Proteomics

Cells were lysed with the PAC lysis buffer (5% SDS, 100 mM Tris pH 8.5, 1 mg/ml chloroacetamide, 1.5 mg/ml Tris (2-carboxyethyl) phosphine), heated at 95 degrees for 30 minutes and then sonicated. Lysate was then randomised and added to a Kingfisher 96 well deep well plate and prepared for an 8h digest protocol on the Kingfisher Duo. Lysate was added to Row G with MagReSyn HILIC beads and acetonitrile is added to 70% final concentration. Rows D, E, and F are filled with 95% acetonitrile and Rows B and C are filled with 70% ethanol. The digest buffer (1 µg/ml MS grade trypsin in 50 mM Triethyammonium bicarbonate) is added to Row A. Desalted peptides were then loaded onto 25cm Aurora Columns (IonOptiks, Australia) using a RSLC nano uHPLC systems connected to a Fusion Lumos mass spectrometer. Peptides were separated by a 70 min linear gradient from 5% to 30% acetonitrile, 0.5% acetic acid. The mass spectrometer was operated in DIA mode, acquiring a MS 350-1650 Da at 120k resolution followed by MS/MS on 45 windows with 0.5 Da overlap (200-2000 Da) at 30 k with a NCE setting of 28. The output was analyzed on DIA-NN (v. 1.8.2 beta 27) with the Mus musculus FASTA. The precursor m/z range was set to 350-1650 and the fragment ion range was set to 200-2000. LFQ intensities for proteins quantified in at least three biological replicates in at least one experimental group were binary-logarithm transformed. Missing values were imputed from a width-compressed, down-shifted Gaussian distribution using Perseus (version 1.6.15.0)^49^.

### Phosphoproteomics

Cells were lysed and proteins were digested as described above. Zr-IMAC beads were then used for phosphopeptides enrichment. The beads were suspended in a bind buffer (0.1M Glycolic acid, 85% acetonitrile and 5% Trifluoroacetic acid) and added to row G of the Kingfisher deep well plates. Another wash with the bind buffer was conducted in row F for equilibration of the beads. The digested peptides were added to bind buffer for a final volume of 200 µl. Two subsequent washes were conducted with 80% acetonitrile, 5% Trifluoroacetic acid, and 0.1M glycolic acid. A final wash was conducted with 10% acetonitrile, 0.2% Trifluoroacetic acid, and 0.1M glycolic acid. Phosphopeptides were eluted in 1% NH4OH, loaded on Evotips and analysed using the Bruker TIMS-ToF SCP coupled with an Evosep LC system on the Whisper 40 SPD method.

### Immunofluorescence microscopy

Coverslips were washed overnight in 1 M hydrochloric acid at 65 °C before being rinsed twice with deionised water and stored in 70% ethanol. Prior to plating cells, coverslips were washed in PBS and coated with 1 μg / mL laminin-I (R&D Systems; #3446-005-01) at 37 °C for 1 hour. 5 × 10^4^ cells were plated on coated coverslips and allowed to adhere for 48 h. Cells were fixed by replacing the media with fixing buffer (3.7% Formaldehyde / 100 mM PIPES / 10 mM EGTA / 1 mM MgCl2 / 0.2% v/v Triton X-100) and incubation at 37 °C for 15 min. Cells were washed once with TBS and incubated in 0.1 M glycine (Sigma-Aldrich; #G8898) in TBS for 10 minutes at room temperature to quench excess formaldehyde before an additional wash with TBS. Permeabilisation was performed with TBS-T for 5 minutes at room temperature. Cells were washed once in TBS supplemented with 0.05% Triton X-100 (v/v) and then blocked with TBS supplemented with 2% BSA and 0.1% Triton X-100 (v/v) for 1 hour at room temperature. Primary antibodies were diluted in blocking buffer and incubated with cells overnight at 4 °C, followed by 3 washes with TBS-T for 5 minutes each at room temperature with gentle agitation. Secondary antibodies and phalloidin-Atto647N (Sigma-Aldrich; #65906, working concentration 25 nM) were diluted in blocking buffer and incubated with cells in the dark for 45 minutes at room temperature. Cells were washed a further 3 times in the dark in TBST at room temperature for 5 minutes each with gentle agitation, followed by a final wash with deionised water. Cells were mounted with ProLong™ Glass Antifade Mountant with NucBlue™ (Invitrogen; #P36981) for nuclear staining. Cells were imaged on Nicon CSU-W1 SoRa microscope using a 405, 488 nm, 568 nm and 647 nm lasers with 100X oil immersion objective. Images were acquired using NIS-Elements AR 5.30.03 (Build 1549) or NIS-Elements AR 6.10.01 (Build 2027) software. To acquire Z-stacks, 4 µm thickness from the cell bottom were imaged with 0.1µm or 0.2µm step size. Images were denoised and deconvolved using NIS-Elements AR 5.21.03 (Build 1489) or NIS-Elements AR 6.10.01 (Build 2027) software and processed using Fiji/ImageJ (v2.14.0/1.54f)^25^. Enhance Contrast was used to help visualize images, except for figure 4A/pMLC2(S19) channel, where the minimum and maximum displayed values were kept the same for all conditions compared.

### MitoTracker staining

MitoTracker deep red FM (Invitrogen; #M22426) was solubilized in DMSO at a stock concentration of 100 μM and stored at −20°C in aliquots. MitoTracker stock solution was diluted in media to a concentration of 400 nM which was added directly to cell culture media already in the culture yielding a final concertation of 200 nM. After a 45 min incubation cells were fixed and permeabilised as described above. To extract mitochondria network features the z-plane closest to the substratum with well-resolved mitochondria, was chosen. Enhance Contrast with 0.1% saturation was used for all conditions, and mitochondria thresholds were generated using Mitochondria analyser (v2.3.1)^24^ in Fiji/ImageJ (v2.14.0/1.54f)^25^ through weighted mean method, 1.25 µm block size and C-value = 9. Mitochondria morphological analysis was performed on the thresholded slices using the same plugin on a field of view basis.

### Immunohistochemistry

Brain samples from mice were fixed in 10% Neutral Buffered Formalin for 16 hours and then transferred to 70% ethanol. Brain samples were then cut into 4 coronal sections before embedding in paraffin blocks. For each brain sample, 8 μm sections were prepared for hematoxilin and eosin (H&E) staining. For immunohistochemistry staining, 8 μm sections were mounted on glass slides. Paraffin removal from sections was achieved by washing samples in xylene for 5 minutes twice. Sections were rehydrated by incubation in decreasing ethanol solutions (100%, 75% and 50% ethanol) for 3 minutes each followed by a rinse in water. Tissue sections were boiled for 5 minutes in 10 mM sodium citrate buffer (0.825 M sodium citrate, 0.175 M citric acid; pH 6.0) for antigen epitope retrieval. Sections were then rinsed in water and washed in TBS-T twice for 5 minutes each. Sections were incubated with Dako REAL peroxidase block solution (Agilent; #S2023) for 5 minutes and rinsed in water. For protein block, sections were treated with serum-free protein block solution (Agilent; #X0909) for 10 minutes. Primary antibodies were prepared in antibody diluent (Agilent; #S3022) and incubated with sections overnight at 4°C. Sections were washed 3 times in TBS-T for 5 minutes each and incubated with DAKO EnVision-HRP rabbit/mouse (Agilent; #K5007) for 45 minutes at room temperature. Sections were washed 3 times in TBS-T for 5 minutes each and developed with DAB-chromagen (Agilent; #K3468) at 1:50 dilution for 10 minutes followed by a rinse in water. Sections were then immersed in Mayeŕs Hematoxylin solution (Agilent; #S3309) for 2 minutes at room temperature and rinsed in water before incubation with Scot’s tap water solution (3.5 g/L sodium bicarbonate, 20 g/L magnesium sulphate) for 2 minutes and a final rinse in water. Sections were dehydrated by incubation in ethanol solutions (50%, 75% and 100% ethanol) for 3 minutes and washed in xylene twice for 5 minutes. Slides were mounted with DPX mountant (Sigma-Aldrich; #06522). Images of stained sections were obtained using a Nanozoomer slide scanner (Hamamatsu Photonics). Ki67 and GFP staining’s were analysed using QuPath software (version 4.0.3). GFP staining was used to distinguish tumour regions.

### Antibodies

Antibodies used for western blotting were anti-FAK (Cell Signaling Technology; #3285, 1:1000), anti-Phospho-FAK (Tyr397) (Cell Signalling Technology; #3283, 1:1000), anti-NF1 (Bethyl; #A300-140A, 1:2000), anti-PTEN (Cell Signalling Technology; #9559, 1:1000), anti-EGFR (Cell Signalling Technology; #4267, 1:1000), anti-COXIV (Cell Signaling Technology; #4850, 1:1000), anti-COFILIN (Cell Signaling Technology; #5175, 1:1000), anti-Glut1 (Cell Signalling Technology;# 73015, 1:1000), anti-Enolase 2 (Cell Signaling Technology; #24330, 1:1000). Primary antibodies used for immunofluorescence microscopy were anti-L-Tubulin (Cell Signalling Technology; #3873, 1:1000), and anti-Myosin Light Chain 2, phospho (Ser19) (Cell Signalling Technology; # 3675, 1:200). Secondary antibodies used for immunofluorescence Alexa Fluor™ 568-conjugated anti-mouse IgG (Invitrogen; #A-11004, all 1:400). Antibodies used for immunohistochemistry were anti-GFP (ChromoTek; # PABG1-20; 1:100) and anti-Ki67 (Cell Signalling Technology; #12202, 1:1000).

### Overall survival of GBM patients

GBM patient survival data was correlated with FAK protein expression (RPPA data) using data from the GBM TCGA database^50^ which was accessed through cBioPortal^51–53^. GBM patients were segregated into two groups according to FAK protein levels. FAK-high and FAK-low groups representing patients with upper (n=50) and lower quartile (n=48) FAK protein expression levels, respectively.

### Computational methods

Where R packages were used for analysis, this was performed using R (v4.4.1). Data were plotted using GraphPad Prism 9 (v9.4.0), R packages; gplots (v3.2.0)^54^, ggplot2^55^, and RColorBrewer^56^. ^13^C_6_-glucose and ^13^C_5_-glutamine isotope tracing data were plotted using Escher-Trace ^57^.

### Quantification and statistical analysis

No statistical methods were used to predetermine the sample size. The investigators were not blinded to allocation during experiments and out outcome assessment. Figure legends contain biological and technical replicate information. Plots include each data point, mean and SD, or mean and SEM. Statistical analyses were performed in GraphPad Prism 9 (v9.4.0) or R (v4.4.1) or Metaboanalyst 5.0^48^. GSEA analysis was performed using the GSEA software (v4.3.1)^58^. *P*-values are represented as ∗*P* < 0.05; ∗∗*P* < 0.01; ∗∗∗*P* < 0.005; ∗∗∗∗*P* < 0.001, between the indicated groups.

## Acknowledgments

We acknowledge Ann Wheeler, Laura Murphy, Martin Lee, James Iremonger and Matthew Pearson of the Advanced Imaging Resource at the Institute of Genetics and Cancer, University of Edinburgh, for their support and expertise. We thank Thomas MacVicar of the The CRUK Scotland Institute, Glasgow, for the insightful discussions. This work was funded by a Cancer Research UK Programme grant award (C157/A24837) to V.G.B. and M.C.F., a joint Brain Tumour Award from Cancer Research UK (C42454/A28596) and The Brain Tumour Charity award (GN-000676) to N.O.C. and M.C.F., and a Cancer Research UK award (A28592) to S.M.P.

## Author contributions

M.C.F. and V.G.B. conceived and co-ordinated the project and helped to design and interpret experiments. R.H.A.M., J.C.D., V.A.G., V.G.B., and M.C.F. designed the experiments. R.H.A.M., J.C.D., V.A.G., A.V.K., J.M.J., G.H., V.G.B., and M.C.F. interpreted the results. R.H.A.M., J.C.D., V.A.G., M.T.M., R.N.C., C.D. performed the experiments. M.C.F., V.G.B., S.M.P., N.O.C. and R.H.A.M. contributed to supervision. R.H.A.M., J.C.D., V.A.G. and C.P. analyzed the data. R.H.A.M prepared figures. R.H.A.M, J.C.D., and M.C.F. wrote the paper. All authors read and approved the manuscript.

---

**Figure S1:**
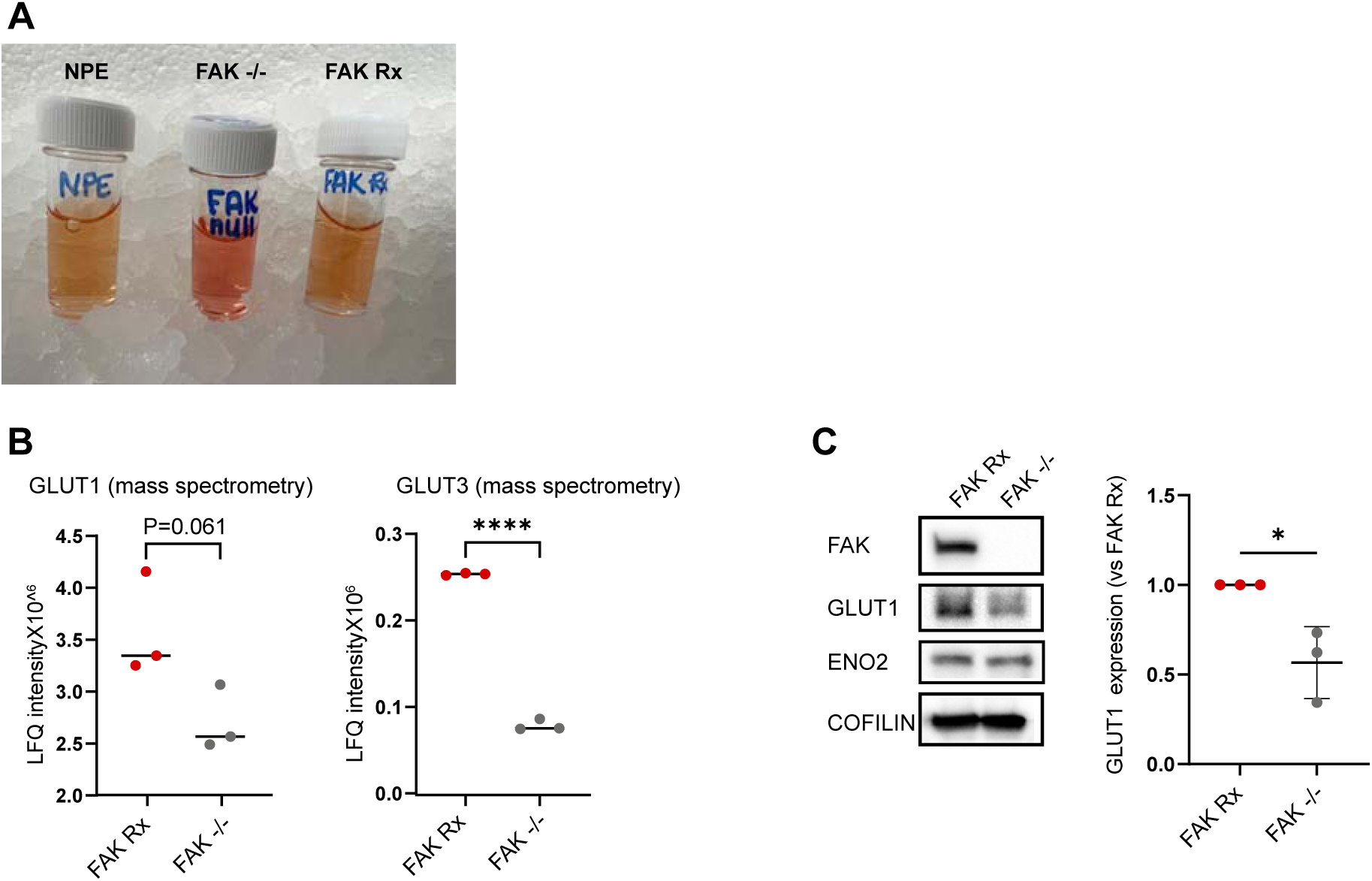
FAK loss is associated with less acidic culture media and reduced Glut1 and Glut3 expression. (A) Representative images of NPE, FAK -/- and FAK Rx culture media, containing the pH indicator phenol red. (B) Label-free quanification (LFQ) mass spectrometrey intensity of GLUT1 and GLUT3 in FAK Rx and FAK -/- cells. Statistics: unpaired two-tailed t-test. (C) Western blot showing FAK, GLUT1, ENO2 expression. COFILIN was used as a loading control. Right panel; quantification of normalised GLUT1 expression from 3 independent experiments. Statistics: unpaired two-tailed t-test.

**Figure S2:**
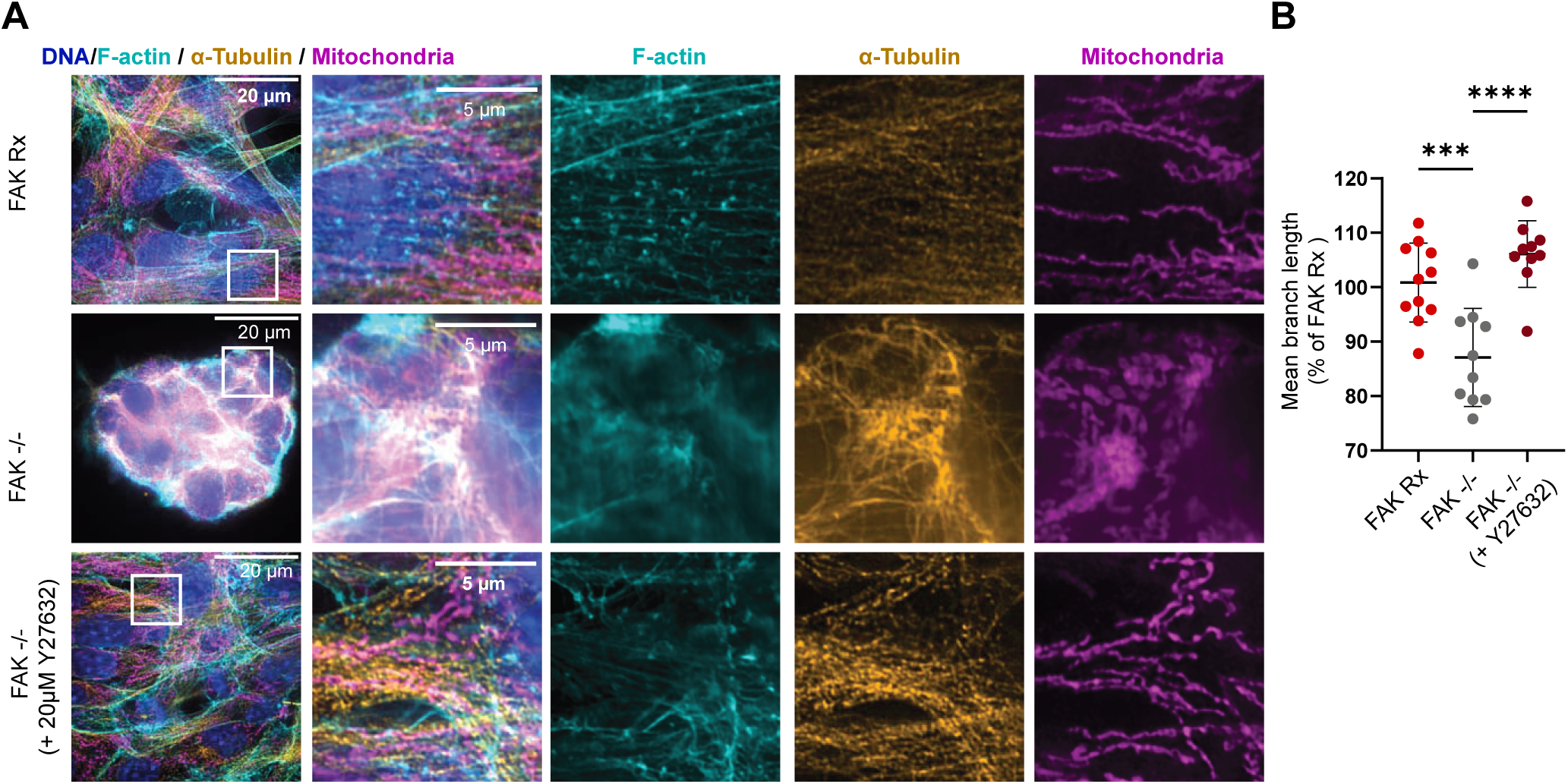
FAK regulates mitochondria morphology through ROCK-pMLC2 S(19) pathway. (A) Representative super-resolution microscopy images for FAK Rx, FAK -/- cells treated with or without 20 µM Y27632 showing mitochondrial morphology stained with MitoTracker Deep Red FM (magenta), the F-actin cytoskeleton labelled with fluorophore-conjugated phalloidin (cyan), α-Tubulin (orange) and nuclei labelled with DAPI (blue). Scale bar is 20 µm in the full field of view and 5 µm in insets. (B) Quantification of mitochondrial mean fragment length in FAK Rx and FAK -/- cells with or without 20 µM Y27632. n = 2 independent experiments. Each dot represents a field of view. Mean and SD are shown. Statistics: one way ANOVA followed by Tukey’s multiple comparisons test.

**Figure S3:**
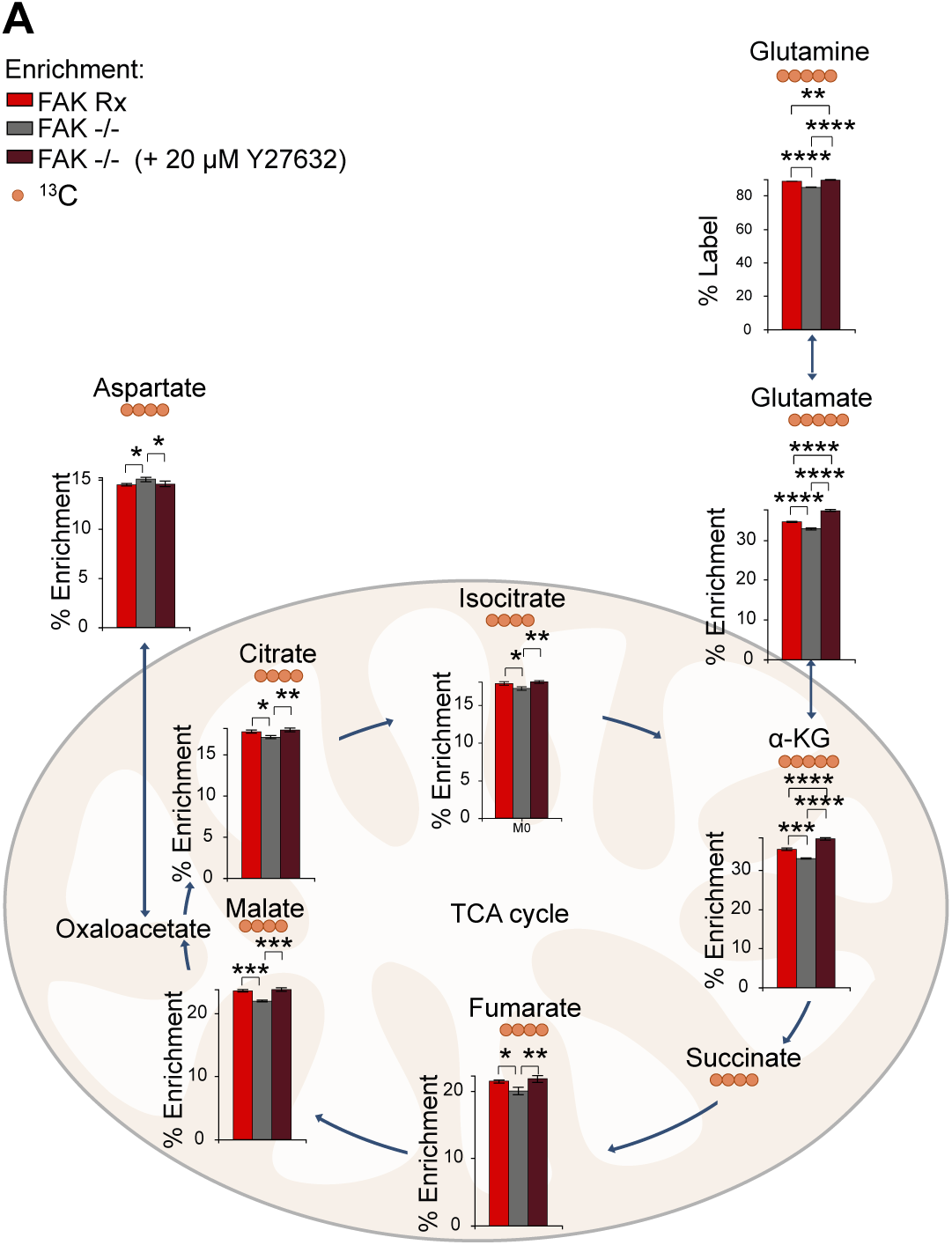
Treating FAK -/- cells with ROCK inhibitor Y27632 increases glutamine oxidation. (A) The atom fraction enrichment of glutamine-derived ^13^C in TCA cycle intermediates after incubation of FAK Rx and FAK -/- cells with^13^C_5_ glutamine for 3h. The main isotopologue of each metabolite is shown and plotted as the fraction of the sum of all isotopologues. Mean and SD are shown. Statistics: one-way ANOVA with a Tukey multiple test correction (n = 3 independent cultures on the same day).

